# Exploring Structural Aspects of the Human Golgi Matrix Protein GRASP55 in Solution

**DOI:** 10.1101/594572

**Authors:** S. Thirupathi Reddy, Luis Felipe Santos Mendes, Natalia Aparecida Fontana, Antonio José Costa-Filho

## Abstract

In mammalian cells, the Golgi apparatus is a central hub for intracellular trafficking, sorting and post-translational modifications of proteins and lipids. The Golgi reassembly and stacking proteins (GRASPs) are somehow involved in the Golgi stacking, which is significant for the proper function of the Golgi apparatus, and also in unconventional protein secretion. However, the structural details on how GRASPs accomplish those tasks are still elusive. In this context, we have explored the biochemical and biophysical properties of the human full-length GRASP55 in solution. Sequence-based analyses and circular dichroism spectroscopy suggest that GRASP55 presents multiple intrinsically disordered sites, although keeping considerable contents of secondary structure. Size exclusion chromatography coupled with multiple-angle light scattering (SEC-MALS) studies show that GRASP55 are monomers in solution. Urea denaturation of GRASP55 suggests that the transition to the unfolded state is a cooperative process. Differential scanning calorimetry (DSC) analysis displays two endothermic transitions for GRASP55, indicating the existence of an intermediate state prior to unfolding. Thioflavin T fluorescence shows that GRASP55 can form protein aggregates/fibrils at the intermediate state. Transmission electron microscopy and fluorescence lifetime imaging microscopy prove that GRASP55 forms large amorphous aggregates but not amyloid-like fibrils in the intermediate state. The significance of these results could be helpful in discussing the proper function of human GRASP55 in the Golgi organization as well as unconventional secretion of proteins.

## 1. Introduction

In mammalian cells, the Golgi apparatus is a central hub for intracellular trafficking and postranslational modifications of proteins and lipids in the secretory pathway [1,2,3]. The Golgi apparatus is formed by laterally linking the stacks of flattened cisternae into ribbon-like structures. This unique architecture is essential for its proper functioning. However, the molecular mechanism underlying how the Golgi stacks of cisternae turn into a ribbon-like structure is still obscure. The Golgi reassembly and stacking proteins (GRASPs) have been reported to play a role in the biogenesis of the Golgi stacks; nonetheless this is still a matter of debate [1–5]. In mammalian cells, GRASP65 and GRASP55 are the two paralogue proteins responsible for the complementary stacking of the Golgi [6]. However, the functions of GRASP proteins are associated but not limited to the Golgi phosphorylation-regulated assembly/disassembly [7,8]. GRASPs are also involved in Golgi remodeling in migrating cells [9], and protein secretion [10]. Furthermore, the cleavage of GRASPs is a necessary event for the fragmentation of the Golgi during apoptosis [11].

GRASP65 and GRASP55 are membrane tethering multitasking proteins, which are distinctively localized to *cis-* and *medial/trans-Golgi* cisternae, respectively. The anchoring between GRASP65 and the cis-Golgi cisternae is established by the interaction with the *cis-* localized golgin GM130 and by the insertion of a myristoyl chain linked to GRASP65 N-terminus [4,12]. GRASP55 predominantly binds to Golgin-45 along with the insertion of its myristoyl or palmitoyl chain into the *medial/trans* Golgi cisternae [13,14]. GRASPs conserve two tandem PDZ domains in their N-termini, which oligomerize into homodimers, thus forming trans-oligomers between the membranes of the cisternae. These *trans*-oligomers are believed to be the “*glue*” to stick the cisternae into a stack [6,15,16] and to connect the Golgi stacks into a ribbon-like structure [12]. In addition, the GRASP C-termini are composed of the more divergent serine and proline-rich (SPR) domain, which is phosphorylated during mitosis to dissociate the *trans*-oligomers with the consequent strip down of the Golgi stacks [2].

Although GRASPs are known to be proteins anchored to the Golgi [4,5], their trans-oligomerization in tethering of the cisternae membranes in solution is not fully understood yet. The crystal structures of various GRASP domains of GRASPs and their complexes with biological partners/ligands have furnished the molecular details to some extent [17–21]. The first crystal structure of human GRASP55 domain, although monomeric, suggested that the possible GRASP55 homodimer might be formed between membranes of cisternae through the interactions of a protruding surface on the PDZ2 domain of one GRASP55 molecule with the binding pocket in the PDZ1 domain of another GRASP55 molecule on the opposite membrane [17].

Feng *et al* [18] proposed, based on the crystal packing symmetry of the rat GRASP domains of both GRASPs (GRASP55 and GRASP65), that GRASP could form dimers through homotypic interactions between the two PDZ2 domains of two neighboring GRASP molecules and the binding of the unstructured C-terminal tail to a pocket in the PDZ1, thus allowing the multimerization. This last work is still very controversial because the GRASP domains do not form dimers in solution and the C-terminal tail observed is not the true in GRASP55/65 C-terminus but rather a truncated construction without the SPR.

A recent report on the structure of the GRASP domain of GRASP55 with the C-terminal peptide of Golgin45 has shown that Golgin45 forms a complex with three PDZ2 domains of different GRASP55 molecules, while PDZ1 domains anchor to the Golgi cisternae membranes and establish a stable cluster of PDZ2 domains, which thus acts as the molecular ‘*glue’* [20]. We can observe that the molecular models obtained so far from the crystal structures are not conclusive on the structural role of PDZ1 and PDZ2 in the formation of GRASPs trans-oligomerization and their interactions with biological partners.

Besides their roles in the membrane tethering, GRASPs have been also implicated in the unconventional secretion of proteins [22–25]. In one of these secretion processes, GRASPs seem to perform a somewhat paradoxical cell function since it involves the bypass of the Golgi apparatus [26]. Kinseth *et al* [27] showed, by gene silencing the GRASP gene in *D. discoideum,* that the cells presented normal growth and protein secretion via conventional mechanisms. However, those cells were not able to form viable sporous, a process that is related to the deficiency in the secretion of the acyl-CoA binding protein (ACBP), which lacks a signal peptide to direct it to the plasma membrane. In a second example, Kmetzsch *et al* [28] reported on the role of GRASP in the unconventional secretion of polysaccharides and virulence factors in the fungus *C. neoformans.* Mutant cells showed alterations in the Golgi morphology and also a reduced capability of secretion of glucuronoxylomannan, one of the major capsular components in *C. neoformans*. Whether the participation of GRASP in unconventional secretion requires its interaction with the plasma membrane is a hypothesis yet to be confirmed. Investigations that aim at exploring GRASP interaction with membranes can contribute to elucidate such issue.

In addition to the various functional properties of GRASPs, it has been recently shown that GRASPs can also exhibit the unusual intrinsically disordered (ID) behavior. Results from our group on GRASPs from both *Cryptococcus neoformans* (CnGRASP) and *Saccharomyces cerevisiae* (ScGRASP, Grh1) revealed the high propensity of ID regions (IDRs) in both the SPR and the GRASP domains [29–31]. Nevertheless, the roles of the intrinsically disordered domains in the functional cycle of both CnGRASP and Grh1 are still unknown. However, whether this ID behavior is a general feature among members of the GRASP family remains to be investigated and leads us to scrutinize the possible ID behavior of human GRASPs.

Towards this goal, as a first step, we have expressed and purified the human full-length GRASP55 and explored its biochemical and biophysical properties in solution. The results on the ID behavior, oligomeric state, chemical unfolding and temperature-dependent aggregation here presented can be useful in understanding the function of human GRASP55 in the Golgi organization as well as unconventional secretion of proteins.

## 2. Materials and methods

### 2.1. Bioinformatics tools

The protein disorder was predicted using the PONDER-FIT [32].

### 2.2. Protein expression and purification

The full-length human GRASP55 gene was synthesized with a codon-optimization for *E. coli* expression (Genscript) and further subcloned in pET28a vector using NDE1 and Xho1 sites. Gene integrity was confirmed by restriction analyses and DNA sequencing. For protein expression, GRASP55-pET28a was transformed in chemically competent *E. coli* Rosetta (DE3). Cells were grown in LB medium supplemented with chloramphenicol (34μg/mL) and kanamycin A (50μg/mL), at 37°C (until reaching an OD_600nm_ of 0.8-1). The gene expression was induced by isopropyl β-D-thiogalactopyranoside (0.5 mM) with overnight incubation (16 h) at 20°C. The cells were harvested, resuspended in lysis buffer (25 mM HEPES, pH 7.4, 300 mM NaCl, 10 mM imidazole, 5 mM β-mercaptoethanol) and then lysed by sonication. The insoluble cell fragments were removed by centrifugation (14,000 × g for 30 min at 4°C). The human full-length GRASP55 was loaded on to the Ni-NTA superflow column (QIAGEN) and then eluted with 300 mM imidazole in lysis buffer, and the flow-through was collected and concentrated by centrifugation. To obtain the pure full-length GRASP55, size exclusion chromatography (SEC) was used to remove the residual impurities by passing through a Superdex 200 10/300 GL gel filtration column (GE Healthcare Life Sciences). The peak fractions were collected and concentrated to 10-12 mg/ml. The purity of human full-length GRASP55 was checked using SDS-PAGE.

### 2.3. Circular dichroism (CD) spectroscopy

The far-UV CD experiments were performed on a Jasco J-815 CD spectrometer (JASCO Corporation, Japan) using a quartz cuvette with a path length of 1 mm at 20°C. Before recording the CD spectra, the instrument parameters were set to a scanning speed of 50 nm/min, spectral bandwidth of 1 nm, a response time of 0.5 sec, and then collected the average spectra from the independent measurements to avoid instability contributions to the final spectra. GRASP55 spectrum was recorded from a sample at 0.1 mg/ml in 20 mM sodium phosphate buffer, pH 8. The CD spectrum was deconvoluted using the CDSSTR software with various databases available at DICHROWEB [33,34]. Urea denaturation experiments were carried out on GRASP55 in 20 mM sodium phosphate buffer, pH 8.0 at 20°C.

### 2.4. Size exclusion chromatography coupled with multi-angle light scattering

Size exclusion chromatography coupled with multi-angle light scattering (SEC-MALS) studies of GRASP55 were performed on a miniDAWN TREOS multi-angle light scattering equipment with detectors at three angles (43.6°, 90° and 136.4°) and a 659 nm laser beam (Wyatt Technology, CA). A Wyatt QELS dynamic light scattering module for determination of hydrodynamic radius and an Optilab T-rEX refractometer (Wyatt Technology) were used in-line with a size exclusion chromatography analytical column (Superdex 200 HR10/300, GE Healthcare) [35]. Prior to any measurement, the samples were centrifuged at 11,000×g for 10 min at 4°C. GRASP55 was eluted through the Superdex 200 column using 50 mM Tris HCl, 300 mM NaCl buffer, pH 8.0 with a flow rate of 0.5 ml/min. Data collection and SEC-MALS analyses were carried out with ASTRA 6.1 software (Wyatt Technology). The refractive index and viscosity of the solvent were defined as 1.331 and 0.890 cP, respectively. dn/dc (refractive index increment) value for all samples was defined as 0.1850 mL/g (a standard value for proteins) [35].

### 2.5. Dynamic light scattering

Dynamic light scattering (DLS) studies of GRASP55 with increasing temperature were carried out using a Zeta-Sizer dynamic light scattering system (Malvern Instruments Ltd, Malvern, UK). The protein sample (2 mg/ml) was subjected to a He-Ne laser of 633 nm and the scattered light measured at a fixed angle of 90°. The measurements were obtained as the mean of three successive counts and carried out in HEPES buffer, pH 7.4 at 20°C using a glass cuvette of 10 mm path length. Prior to any measurement, the sample was centrifuged at 11, 000×g for 5 min at 4°C.

### 2.6. Differential scanning calorimetry

Differential scanning calorimetry (DSC) studies of human GRASP55 were performed on a Nano-DSC II–Calorimetry Sciences Corporation, CSC (Lindon, Utah, USA). Protein solutions at 1.5-2.5 mg/ml were used in HEPES buffer, pH 7.4. Before running the DSC scan, the samples were degassed for 15 min utilizing high vacuum desiccation to avoid bubbles in the solution and then used to fill the sample cell. Besides, the reference cell was filled with HEPES buffer, pH 7.4 without protein. The DSC scan was recorded from 0 to 100°C with a scan rate of 0.5°C/min. The protein sample was subjected to two heating and one cooling scans and then, the first heating scan was used for further analysis after the blank subtraction, baseline correction, and normalization. The protein unfolding or melting temperatures were measured by considering the peak maximum of the transition peaks and the corresponding enthalpies were calculated by integrating the area under the transition peaks [36,37].

### 2.7. Fluorescence spectroscopy

Steady-state fluorescence measurements were performed using a Hitachi F-7000 equipment. The excitation and emission slit widths of 5 nm each were used in all the experiments. The chemical unfolding studies of GRASP55 (10 μM) were carried out using the tryptophan fluorescence by exciting at 295 nm with increasing concentration (0 to 8 M) of urea and the corresponding emission spectra were recorded from 305 to 400 nm. Background intensities of all samples were subtracted from each spectrum to avoid any contribution due to solvent Raman peak and any other scattering artifacts. Besides, the tryptophan anisotropy of GRASP55 (10 μM) was measured with increasing concentration (0 to 8 M) of urea. The effect of temperature on the oligomeric state of GRASP55 was explored using Thioflavin (ThT) fluorescence. In this case, solutions of ThT at 15 μM and protein at 30 μM were used. The same experimental parameters described above were used to record the fluorescence spectra of ThT.

### 2.8. Transmission electron microscopy

Transmission electron microscopy (TEM) experiments were carried out with the native (20°C) and pre-heated (to 60°C) samples of GRASP55 using a JEOL JEM 100 CXII. To record the TEM images, a drop (2 μL) of the native and pre-heated to 60°C (intermediate state) protein samples (30 μM) were placed on the copper grids separately and then incubated for 5 min at room temperature. The samples were then stained with 2% uranyl acetate for 5 min and followed by removal of the excess aqueous medium by blotting with a filter paper.

The TEM images were collected at room temperature with the acceleration voltage of 100 kV. The size of protein aggregates was measured using ImageJ software [38].

## 3. Results and discussion

### 3.1. Characterization of native human full-length GRASP55 in solution

The human full-length GRASP55 was successfully purified as described above (Figure 1a). The Far-UV CD spectrum (190-260 nm) of GRASP55 recorded at 20°C is shown in Fig. 1b. The spectrum showed that the GRASP55 exhibited broad bands at ~203 and ~222 nm, suggesting the presence of the disordered as well as ordered secondary structure elements in the protein. The CD spectrum was further analyzed via CDSSTR routine using a set of various databases at Dichroweb server [33,34]. The results from CDSSTR deconvolution showed that full-length GRASP55 has ~21% of α-helix, ~20% of β-sheet, ~17% of turns, and ~42% of disordered regions with the normalized root mean square deviation (NRMSD) of 0.19, which indicates, as already mentioned above, the presence of multiple disorder sites alongside well-structured regions.

**Figure 1.**
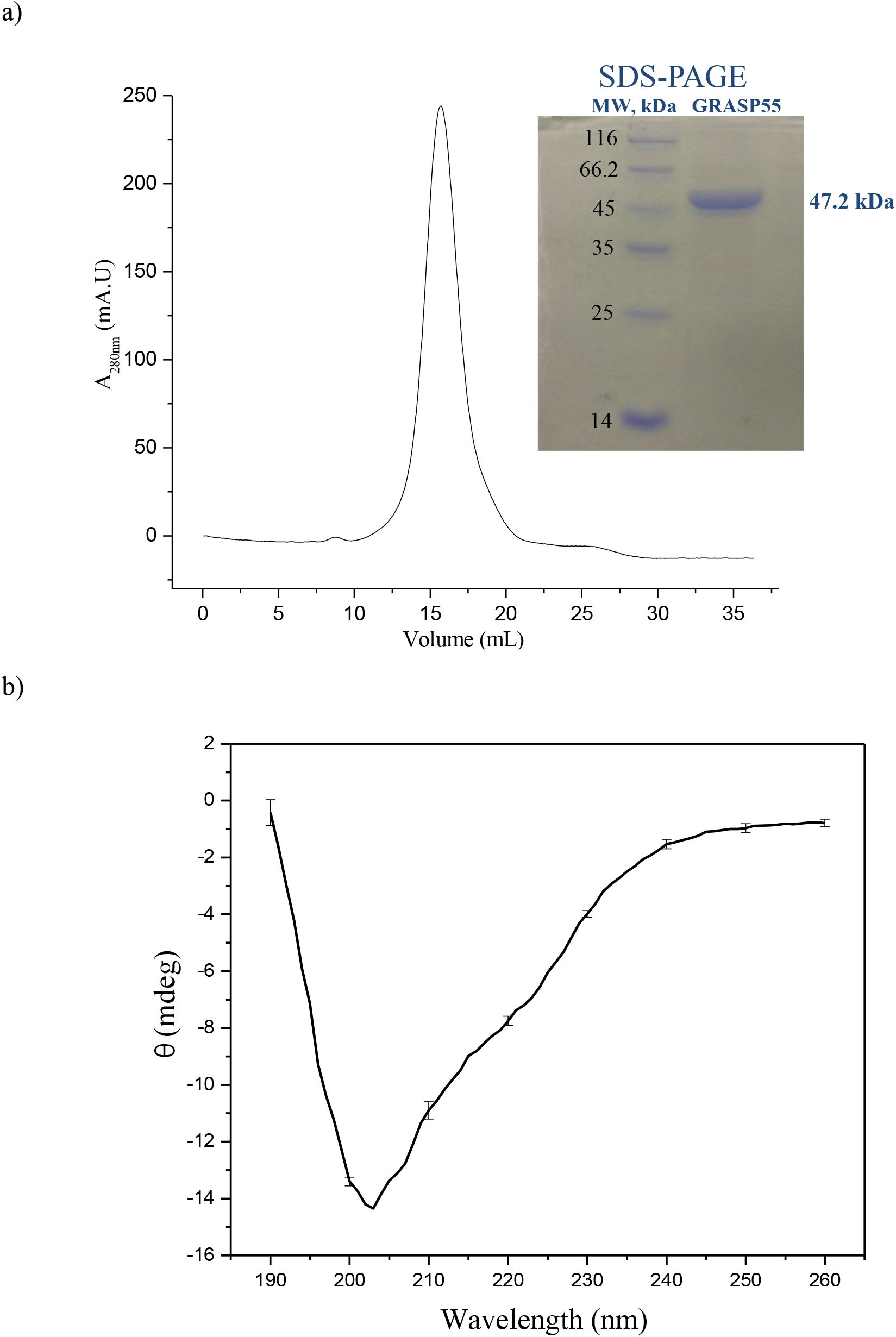
(a) The size exclusion chromatography profile of the GRASP55. Inset shows the gel picture obtained from the SDS-PAGE indicating the purity of GRASP55. (b) The far-UV CD spectrum of full-length GRASP55 in 20mM phosphate buffer, pH 8.0 at room temperature.

In addition, the protein family database analysis showed that the full-length GRASP55 consists, as expected, of two domains: the GRASP (residues 1-208) [17], including the two PDZ subdomains, and the SPR (residues 209-452). Furthermore, the sequence-based prediction of protein disorder suggested that the C-terminal (SPR) domain of full-length GRASP55 has high propensity of intrinsic disorder (Figure 2), which is similar to the SPR domains of the CnGRASP and ScGRASP [29,30].

**Figure 2.**
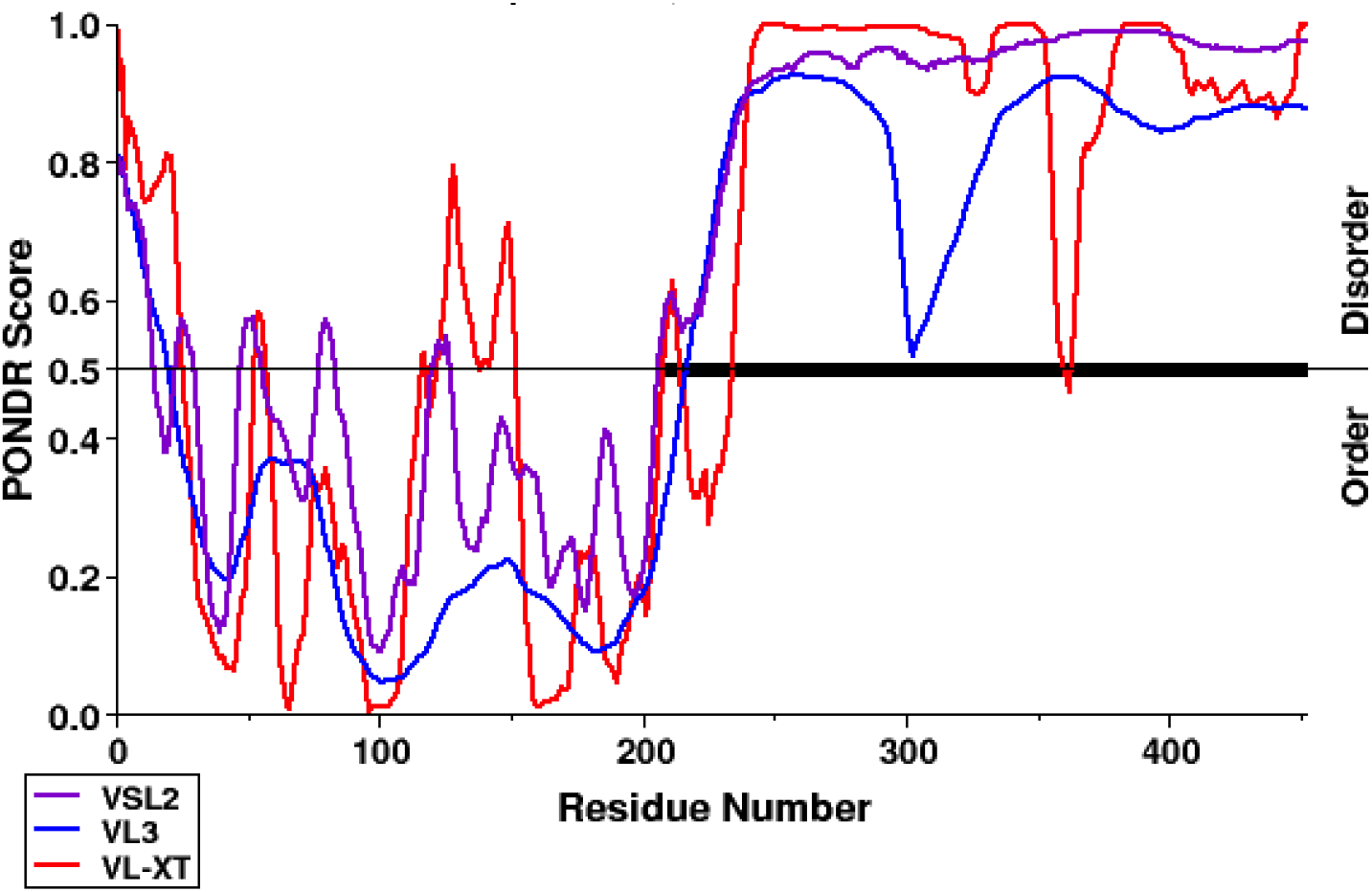
Disorder prediction of GRASP55 using VSL2 (violet), VL3 (blue) and VL-XT (red) programs. The black line in the middle indicates the limit above which the residue has more than 50% propensity of residue disorder.

The molecular mass of full-length GRASP55 was estimated to be 47.2 kDa using ProtParam tool [39], which agrees with the SDS-PAGE result in Figure 1a. In order to determine the oligomeric state of GRASP55 in the native state, SEC-MALS was employed to estimate the accurate molar mass (MM) and the hydrodynamic radius (Rh) of the protein in solution without the need to use globular proteins as standards for mass determination (Figure 3). The SEC-MALS data showed low polydispersity index and yielded values of MM and R_h_ of 46.7 kDa and 4.3 nm, respectively. The obtained MM of the protein clearly indicates that GRASP55 exists as a monomer in solution. Furthermore, the obtained R_h_ value of GRASP55 is much higher than what is expected for a ~47 kDa protein (~3 nm) [40,41] based on the relation between Rh and molecular mass of different standard proteins. This is another indication that GRASP55 exhibits intrinsically disorder (ID) behavior in solution and agrees with our findings mentioned above. The ID behavior previously observed for lower eukaryotic GRASPs [29,30] thus seems to be a general property of GRASPs, spanning from lower to higher eukaryotes.

**Figure 3.**
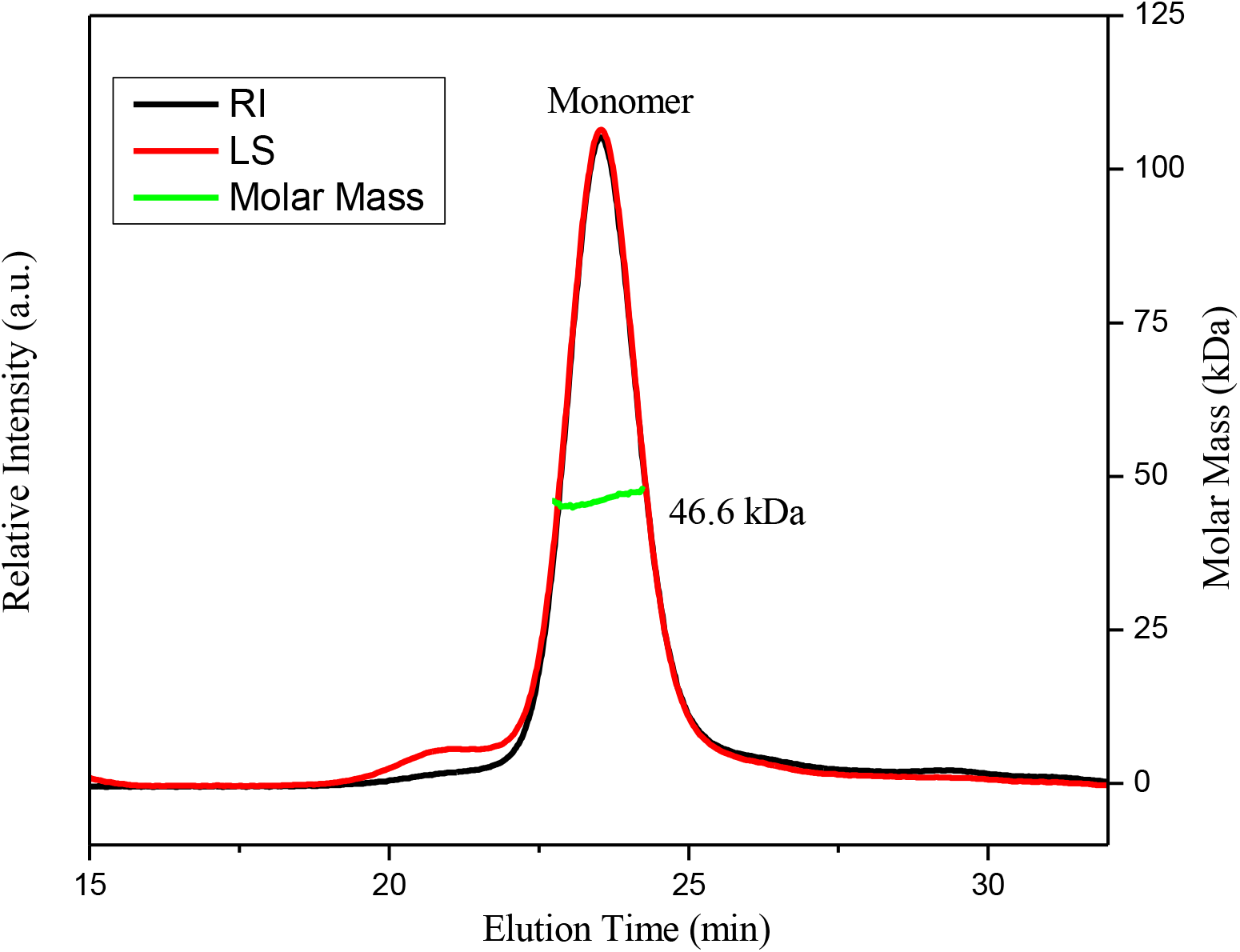
SEC-MALS of GRASP55. The chromatogram display the light scattering (LS) at 90° angle and refractive index (RI) curves together with the molar mass (MM) of the peak calculated by MALS.

### 3.2. Chemically-induced unfolding of GRASP55

Chemical unfolding of GRASP55 was monitored by measuring the intrinsic tryptophan fluorescence in the presence of increasing concentrations of urea (up to 8 M). The fluorescence emission spectra of GRASP55 at different concentrations of urea are shown in Figure 4a. The emission maximum (λ_max_) of GRASP55 in native conditions was centered at 334 nm, which agrees with the location of the tryptophans (W90, W103, and W184) of GRASP55 in the more ordered (N-terminal) region of the protein as shown in the crystal structure of GRASP55 GRASP domain [17]. The λ_max_ exhibited the red shift (increase in λ_max_) with the addition of urea (up to 3 M) to the protein solution (see inset in Figure 4a). This indicates that GRASP55 gradually unfolds its native structure and also suggests the tryptophans in the ordered region are slowly exposed to the solvent. A sudden increase in the λ_max_ was noticed at ~4 M (transition midpoint) urea concentration. Subsequently, at higher concentration (>5 M) of urea, the λ_max_ of the GRASP55 remains constant (352 nm) despite the further increase of urea (up to 8 M). Moreover, the fluorescence anisotropy of GRASP55 also showed a sudden change (decrease) in the tryptophan anisotropy at ~4 M (transition midpoint) urea concentration (Figure 4b), which agrees with the abrupt change (increase) in the emission maximum of tryptophan fluorescence.

**Figure 4.**
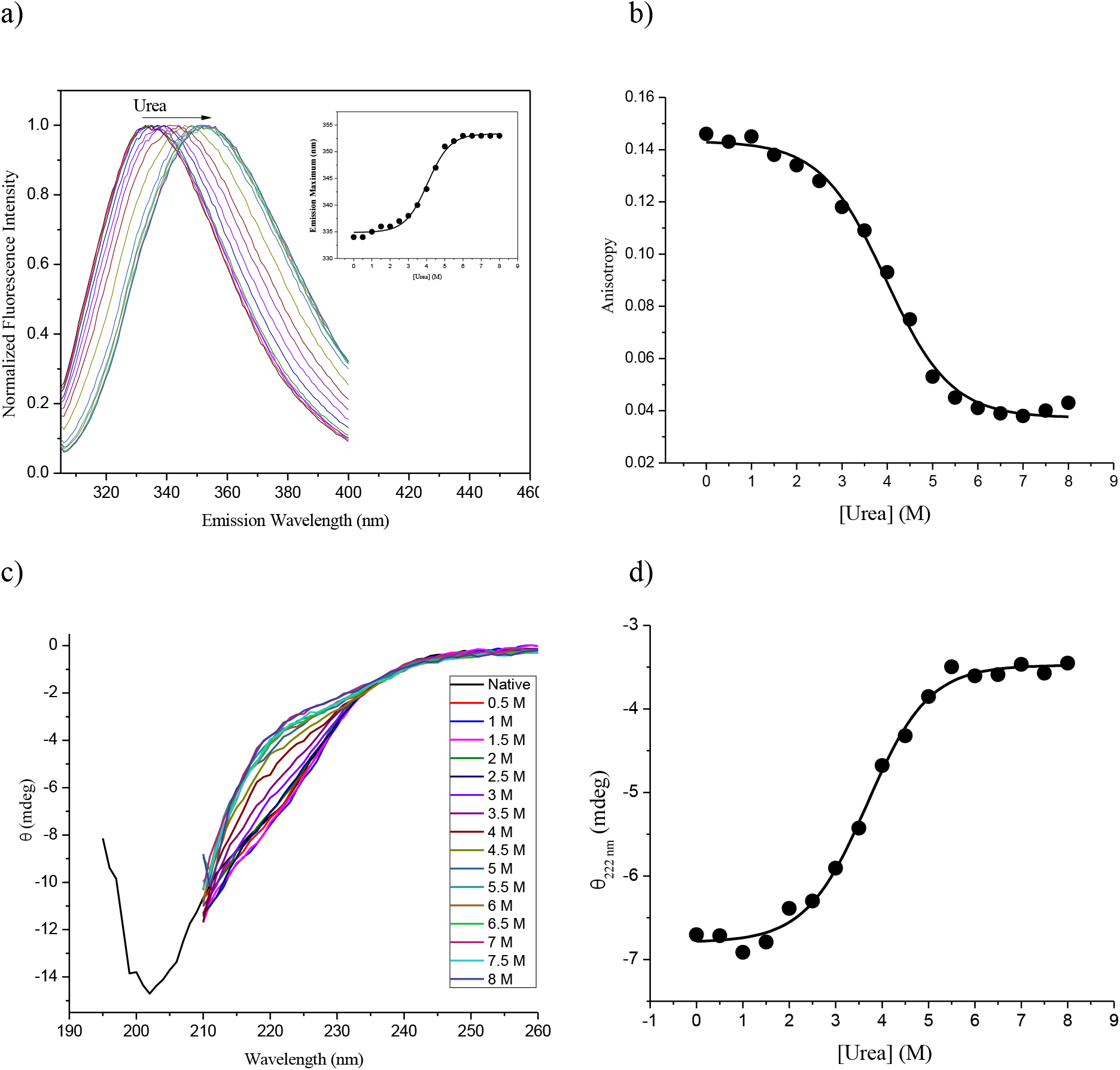
(a) The tryptophan fluorescence spectra of GRASP55 with increasing concentration of urea. The inset shows a plot between the emission maximum of GRASP55 with increasing concentration of urea. (b) A plot of tryptophan anisotropy of GRASP55 with increase in concentrations of urea. (c) Far-UV CD spectra of GRASP55 with increasing concentration of urea. (d) A plot of ellipticity (mdeg) at 222 nm with increase in the concentrations of urea.

In addition, the urea-induced denaturation of GRASP55 was also investigated using CD spectroscopy. The CD spectra of GRASP55 with increasing concentration of urea are displayed in Figure 4c. The variation in the secondary structure of GRASP55 was followed by monitoring the ellipticity at 222 nm. Figure 4d shows that the ellipticity of GRASP55 decreases gradually with the addition of urea, which indicates the loss of secondary structure of the protein. The transition midpoint in the ellipticity of the protein is again at ~4 M urea concentration, which suggests that the exposure of the Trp residues happens concomitantly with the loss of secondary structure. With further increase in urea (5.5 M and above) concentration, the secondary structure of the protein is completely lost and the ellipticity at 222 nm remains constant (up to 8 M). Therefore, the secondary structure of GRASP55 gradually decreases with increasing concentrations of urea and then unfolds completely at higher concentration of urea. The somewhat high value of the transition midpoint (ca. 4 M urea), observed both from fluorescence and CD data, suggests that the monomeric GRASP55 presents a very stable structure in the native state.

### 3.3. Temperature-induced unfolding of GRASP55

To broaden our studies concerning the structural behavior of the GRASP55 monomer in solution, we also examined the stability of the GRASP55 monomer upon variations in temperature. In this direction, the thermotropic behavior of GRASP55 was firstly investigated utilizing differential scanning calorimetry (DSC) (Figure 5a). DSC data on native GRASP55 exhibited two endothermic transitions centered at ~50 and ~74°C. The first transition could be attributed to the monomers (native state) turning into higher ordered oligomers or aggregates (intermediate state), which would then dissociate and giving rise to the second peak.

**Figure 5.**
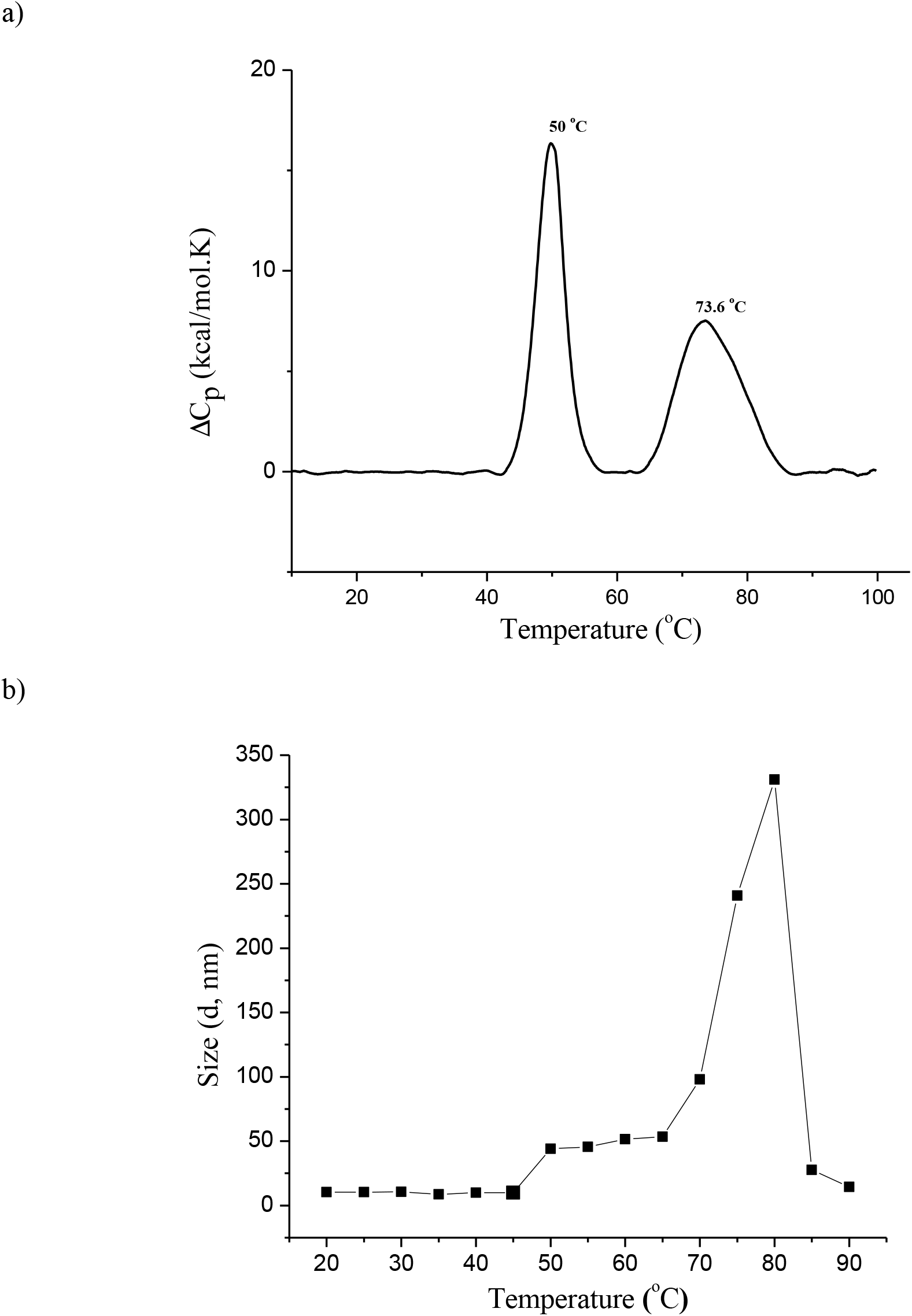
(a) The DSC thermogram (0 to 100°C) and (b) effect of temperature on the size of GRASP55 in HEPES buffer pH 7.4.

The transition enthalpies of the first and second peaks were calculated by integrating the area under the transition curves and the obtained values were 108 and 128 kcal/mol, respectively. The transition enthalpy of the second transition is higher than the first one and it indicates the thermal unfolding of GRASP55 from the intermediate state (higher ordered oligomers/aggregates) is more energetic than the transition of the native to intermediate state. When the GRASP55 sample was subjected to a second heating scan right after the first cooling scan, there were no transition peaks observed in the thermogram (not shown). This clearly indicates that the overall thermal unfolding of GRASP55 is an irreversible process.

To check the thermal reversibility of the first transition in the DSC thermogram (monomers to intermediate state), the protein sample was heated to 60°C (intermediate state) in the first heating scan with a scan rate of 0.5°C/min and subsequently cooled to 0°C, and then reheated to 60°C. The first heating thermogram (Figure S1) clearly shows that GRASP55 exhibited the same sharp transition peak at ~50°C seen in Figure 5a, which corresponds to the transition of monomers (native state) to the intermediate state. After cooling, in the second heating scan, the sample was reheated to 60°C but then no peak was observed (below 60°C) (Figure S1). This clearly indicates that the temperature-induced transition of monomers of GRASP55 to intermediate state is also a thermally irreversible process.

The DSC data suggested that with increasing temperature GRASP55 would form either higher order oligomers or aggregates from monomers in the native state. To examine the effect of temperature on the size of GRASP55, we have performed DLS experiments at different temperatures (from 20 to 90°C) with a step size of 5°C and the results are shown in Figure 5b. At initial temperature (20°C), GRASP55 exhibited the size (diameter) of ~10 nm that remained the same up to 45°C, which indicates that the native GRASP55 is thermally stable up to 45°C as seen in the DSC thermogram (Figure 5a). At 50°C, an increase to ~50 nm in the size of GRASP55 was observed, which suggests that the first peak in the thermogram would be due to the transition of monomers to higher order oligomers or aggregates. From 50 to 65°C, the size of GRASP55 remained almost constant, indicating the maintenance of the population of higher order oligomers or aggregates with temperature. A sharp increase in the size of the protein was noticed from 70 to 80°C and then an abrupt decrease after 80°C, reaching the size of ~10 nm at 90°C. This clearly shows that the second broad peak in the GRASP55 thermogram can be attributed to the unfolding or dissociation of protein higher order oligomers or aggregates.

Our group has previously reported the formation of fibril-like structures by Grh1, the GRASP from *S. cerevisae,* induced by temperature and other conditions [30]. Hence, to explore the nature of the intermediate state of GRASP55 in more detail and whether or not GRASP55 was also forming fibril-like structures, we used the ThT fluorophore that has been shown to bind to hydrophobic sites in protein structures in solution [42–45]. The maximum of fluorescence intensity of ThT centered at 485 nm is decreased as the protein sample is heated from 20 to 45°C (Figure 6a), which indicates that the ThT bound to GRASP55 monomers is gradually released due to the conformational changes of the protein with increasing temperature. Thereafter, the fluorescence intensity of ThT abruptly increased (Figure 6b), suggesting the transformation of protein monomers into either aggregated or fibrillated structures [45] for which ThT has higher affinity. This structural transition of monomers to protein aggregates/fibrils at ~50°C (Figure 6c) exactly matches the first transition peak in our DSC thermogram of GRASP55 (Figure 5a).

**Figure 6.**
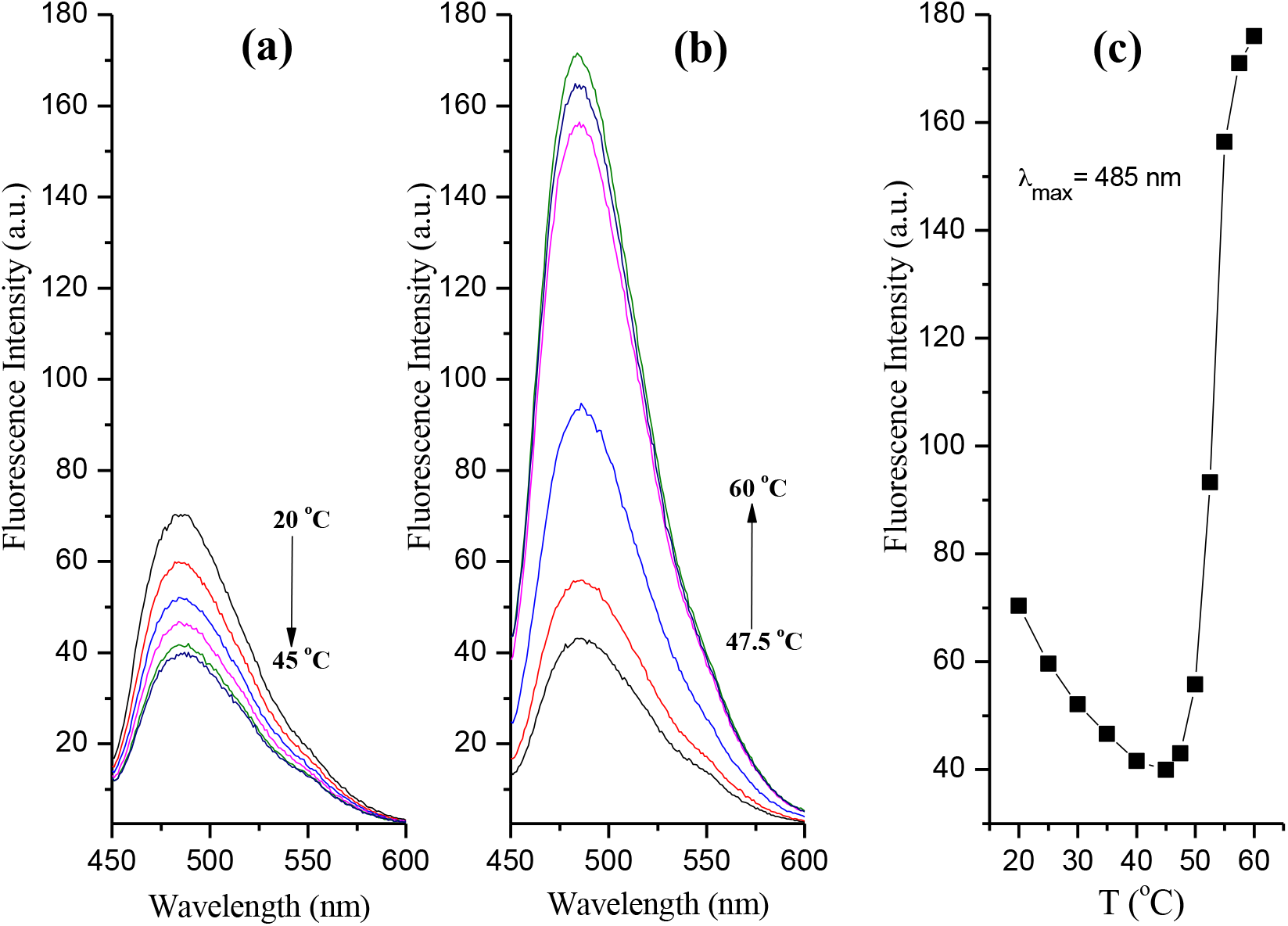
The fluorescence spectra of ThT in the presence of GRASP55 with increasing temperature (a) from 20 to 45°C and (b) 47.5 to 60°C. (c) A plot of emission maximum of ThT at 485 nm in presence of GRASP55 with increasing temperature.

To further investigate the thermal stability of GRASP55 aggregates/fibrils formed in the intermediate state, the protein sample heated to 60°C (intermediate state) was allowed to cool to room temperature and then, at this temperature, the fluorescence of ThT measured again. In this case, ThT fluorescence exhibited the highest intensity (Figure S2), which indicates high propensity of binding this fluorophore by the aggregates/fibrils. Subsequently, with increasing temperature of pre-heated protein sample, the ThT fluorescence exhibited a gradual decrease in its intensity (Figure S2). This observation clearly suggests the disassembly of the protein aggregates/fibrils with increasing temperature. These results also explain why we did not observe the first transition peak in the second heating scan of GRASP55 with increasing temperature (Figure S1).

To evaluate the structures formed by the GRASP55 with increasing temperature, transmission electron microscopy (TEM) studies of the native and pre-heated (60°C) GRASP55 were performed at room temperature (20°C) and the results are shown in Fig. 7. The TEM image of GRASP55 in the native state (Figure 7a) showed small spherical particles with the mean size of 12±3 nm. On the other hand, in the intermediate state (60°C), GRASP55 forms larger amorphous aggregates of different sizes (up to ~1 μm in length) in solution (Figure 7b). Therefore, the TEM measurements suggest that the GRASP55 forms larger aggregates in the intermediate (60°C) state (see Figure 7b) but not fibrils.

**Figure 7.**
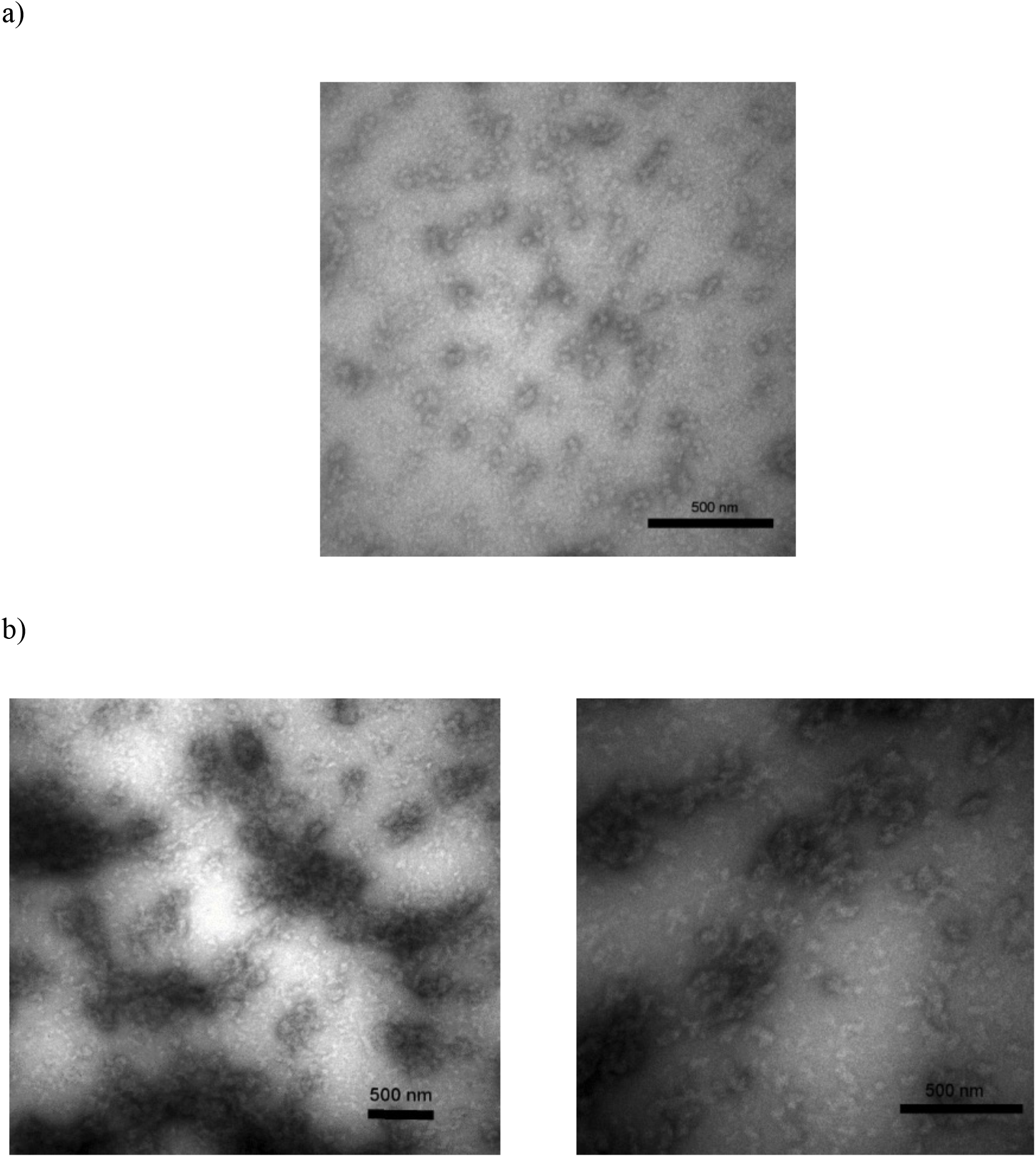
TEM images of (a) native and (b) pre-heated (to 60°C) samples of GRASP55 recorded at room temperature (20°C).

The aggregation of GRASP55 at the intermediate state was further investigated by the fluorescence lifetime imaging microscopy (FLIM) (Figure S3). FLIM allows one to specifically monitor fibrillation based on the intrinsic fluorescence in the visible range arising from the delocalization of electrons between the peptide backbones in the fibril β-sheet structures [46,47]. Our recent report [30] suggests that the *S. cerevisae* GRASP, Grh1, forms fibrils upon heating by the emission of fluorescence in the visible region upon excitation with UV light [48]. Here, we did not see any fluorescence in both the native and intermediate states (Figure S3), which suggests GRASP55 forms protein aggregates but not amyloid-like fibrils in the intermediate state.

## 4. Discussions

GRASP55 is one of the GRASP family members in mammals, which are the only organisms possessing two GRASPs paralogue genes. Although several studies on the roles played by GRASP55 in the cell have been reported [5,6,25,49,50], the details of what the molecular determinants of a specific function would be are still not fully understood. Structural data of the GRASP domain only have shown that even the molecular mechanism of trans-oligomerization in tethering of membranes is still open for debate [17–21]. Moreover, our group has recently shown that although the GRASP domains of GRASP55 and GRASP65 share a common 3D folding, based on the crystallographic data, their behavior in solution can be quite different, making GRASP65 closer to GRASPs from lower Eukaryotes than GRASP55 [31]. Why then mammals would need two GRASP paralogue genes, instead of only one as observed in all other organisms, is an issue that deserves more attention.

It has been recently shown that GRASP55 and the unfolded protein response (UPR) pathway are involved in the control of the unconventional secretion of interleukin 1β in lipopolysaccharide-activated macrophages [51]. The transition from a dimer when in the Golgi membrane to the monomeric state of GRASP55 can then be relevant to control its participation in unconventional protein secretion (UPS). Furthermore, it has been also shown that endoplasmic reticulum (ER) stress induces a strong reduction in the level of GRASP55 dimers [49]. It seems thus clear that new pieces of information on how GRASPs respond to stress-like conditions can help understanding their participation in those processes.

Here we explored full-length GRASP55 biophysical properties by forcing structural transitions upon variations of two parameters of the protein’s environment: urea concentration and temperature.

We showed that, unlike CnGRASP, full-length GRASP55 is a monomer in solution and presents ID behavior more restricted to its SPR domain, which resulted in more cooperative urea-induced unfolding transitions monitored by the intrinsic fluorescence of the Trp residues in the GRASP domain. The intrinsic fluorescence of Trp also indicated the GRASP domain of GRASP55 is somewhat more compact, with wavelength of maximum emission around 334 nm, when compared to other GRASPs, whose maximum emission occurs at wavelength greater than 342 nm [29,30]. As previously observed for other GRASPs [29,30], the overall protein structure seems to be very stable with a transition midpoint around 4 M of urea.

In the thermal unfolding of GRASP55, we showed the protein presented a two-step process. Unlike Grh1, which can form fibril-like structures, GRASP55 forms non-fibril aggregates as intermediate state of the thermal unfolding. GRASP55 has a long intrinsically-disordered SPR domain, which corresponds to more than half of the protein’s residues. This long disordered region might be responsible for the formation of the amorphous aggregates, which differ from the more organized fibril-like structures likely resulting from the organization of the considerably shorter SPR domain of Grh1. The involvement of ID regions in the formation of higher-order oligomers of proteins has been suggested [52] and the final arrangement of such oligomers might be dependent, in the case of GRASPs, on the size and on the number of the specific ID region. It is of course obvious that the temperature needed for the transition of GRASP55 to the amorphous aggregates (ca. 60°C) is not physiological, but it can still be thought of as a way to unravel structural properties of GRASP55 that could be triggered by other stress condition found during the cell life cycle.

The fact that full-length GRASP55 and Grh1 form different types of aggregates (amorphous and fibril-like, respectively) when subjected to stress-like conditions can be related to both the size of their SPR domains and to the number of IDRs within the protein structure. This variety of possible structural arrangements gives rise to the broad structural plasticity of GRASPs, which is likely involved in the plethora of roles that has been attributed to GRASPs over the last few years [1,2].

The different conditions used (urea or temperature) to search the conformational space of GRASP55 led to distinct final structural states. This suggests that different stress inducers could lead to different structural arrangements of GRASP55. A specific stress condition could thus constitute the trigger for a determined structural transition that would bring the same protein to different structural states, which would then act according to the needed function.

## 5. Conclusion

In order to explore the structural properties of human GRASPs, in this study, we performed a broad biochemical and biophysical investigation on the human full-length GRASP55 in solution. The protein disorder prediction tools and CD spectroscopy suggest that full-length GRASP55 show ID behavior in solution. In this case and unlike other GRASPs, ID behavior is more restricted to the protein’s SPR domain. The SEC-MALS data suggest that GRASP55 forms monomers in solution in native conditions with R_h_ value higher than what would be expected if GRASP55 was a regular globular protein. Urea denaturation studies show that GRASP55 has a stable form at native conditions and unfolds at high concentration of urea. The thermal stability of GRASP55 was investigated by DSC and exhibited an intermediate form prior to unfolding. The results of ThT fluorescence suggested that GRASP55 forms amorphous aggregates/fibrils in a temperature-dependent manner. TEM and FLIM studies proved that the GRASP55 forms large amorphous aggregates of different sizes at the intermediate (60°C) state. The results obtained in the present study could be useful in understanding how GRASPs can accomplish the broad spectrum of functions they have been related.

## Abbreviations

CD: Circular dichroism;
DSC: Differential scanning calorimetry;
DLS: Dynamic light scattering;
GRASPs: Golgi-reassembly and stacking proteins;
ID: Intrinsic disorder;
SEC-MALS: Size exclusion chromatography coupled with multiple-angle light scattering;
ThT: Thioflavin T;
TEM: Transmission electron microscopy

## Acknowledgments

The authors thank the Brazilian agencies Conselho Nacional de Desenvolvimento Científico e Tecnológico (CNPq) and Fundação de Amparo à Pesquisa do Estado de São Paulo (FAPESP) for the financial support through Grants No. 2015/50366-7, 2012/20367-3, and 308380/2013-4. STR, LFSM, and NAF thank FAPESP for the post-doctoral and PhD fellowships (Grants No. 2017/12146-0, 2017/24669-8, and 2016/23863-2, respectively). The authors also thank the “Grupo de Biofísica Molecular Sérgio Mascarenhas” from the Physics Institute of University of São Paulo for the access to the SEC-MALS (FAPESP Grant No. 15/16812-0) and DLS instruments, and acknowledge the support of Dra Andressa Patricia Alves Pinto.

