## Supplemental figures for "Exploring Structural Aspects of the Human Golgi Matrix Protein GRASP55 in Solution"

### 1. Methods

#### *1.1. Fluorescence lifetime imaging microscopy*

Fluorescence lifetime imaging microscopy (FILM) was used to investigate the aggregation behavior of full-length GRASP55 on a PicoQuant MT 200 microscope at room temperature. For that, native and pre-heated (to 60 °C) samples of GRASP55 were utilized to record FLIM images. Excitation and emission wavelengths used were of 375 and 405 nm, respectively.

#### Supplementary Figures

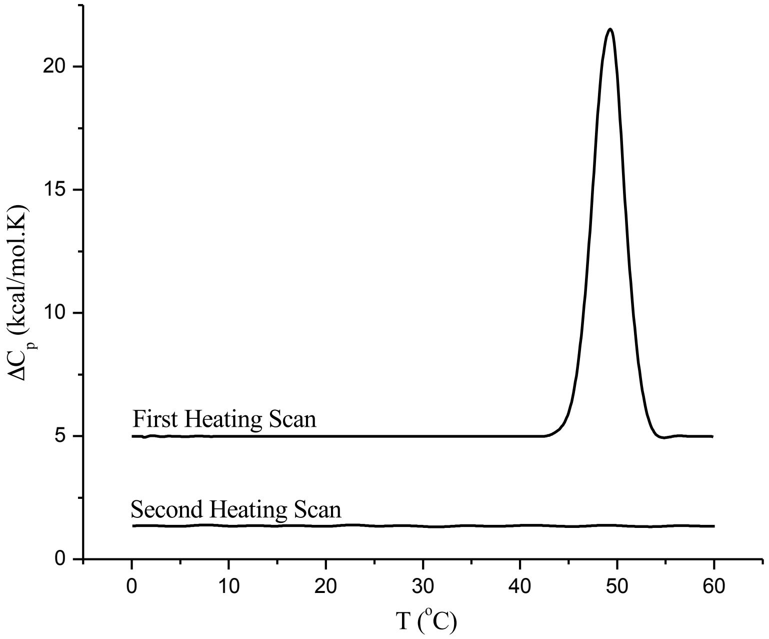


**Figure S1.** The DSC thermograms (0 to 60 °C) of GRASP55 in HEPES buffer pH 7.4.

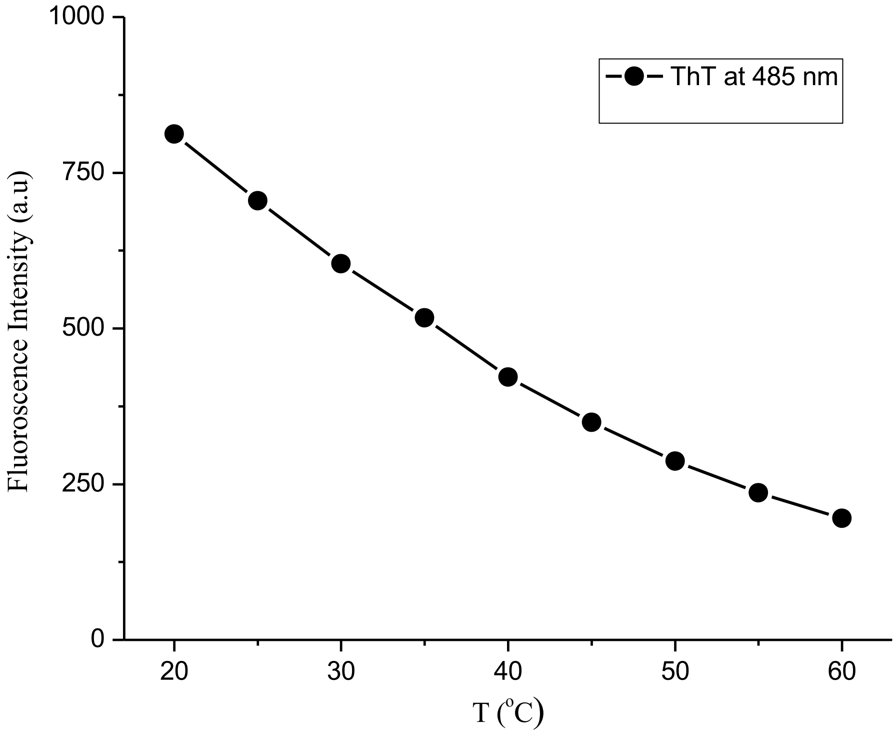


**Figure S2.** Effect of temperature on the ThT fluorescence in the presence of pre-heated (to 60 °C) GRASP55.

(a)

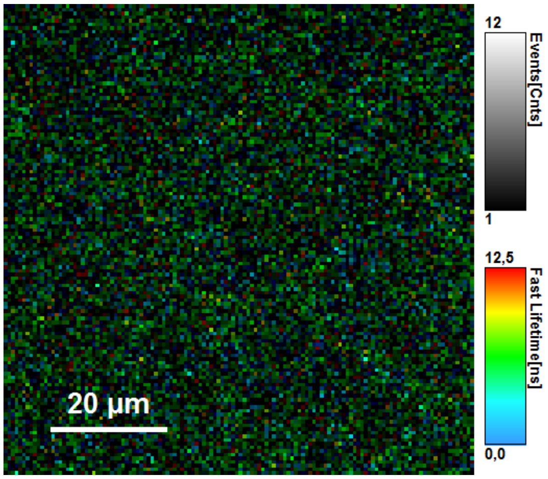


(b)

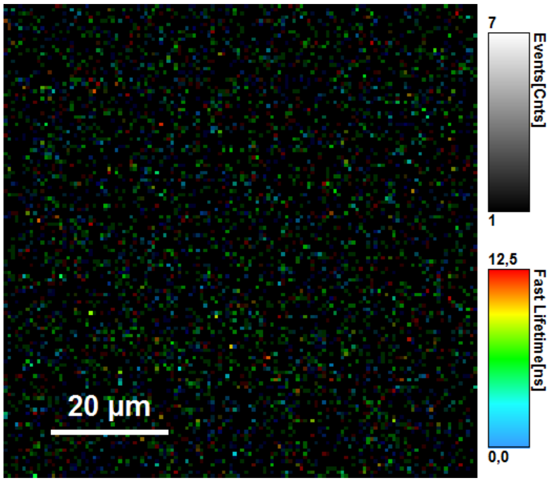


**Figure S3.** The fluorescence lifetime imaging microscopy (FLIM) images of (a) native (20 $^{o}$C) and (b) pre-heated (to 60 $^{o}$C) GRASP55 samples were recorded at room temperature.
